# Rapid method for generating designer algal mitochondrial genomes

**DOI:** 10.1101/2019.12.19.882662

**Authors:** Ryan R. Cochrane, Stephanie L. Brumwell, Maximillian P.M. Soltysiak, Samir Hamadache, Jennifer G. Davis, Jiayi Wang, Samuel Q. Tholl, Preetam Janakirama, David R. Edgell, Bogumil J. Karas.

**Affiliations:** Department of Biochemistry, Schulich School of Medicine and Dentistry, The University of Western Ontario, London, N6A 5C1, Ontario, Canada; Designer Microbes Inc., London, Ontario, Canada

**Keywords:** Mitochondria, Synthetic Organelles, Transformation-Associated Recombination Cloning, Diatom, Eukaryotic Algae, Whole Genome Cloning, *Phaeodactylum tricornutum*

## Abstract

With synthetic biology, we can turn algae into bio-factories that produce high-value molecules (e.g. medicines or biofuels) or tackle global challenges (e.g. malnutrition and climate change). This realization has provoked rapid progress towards the creation of genetic tools for multiple algal species, notably *Phaeodactylum tricornutum.* The power of synthetic biology to generate more useful or productive organisms is contingent on the ability to produce diverse DNA molecules and rapidly screen them for beneficial variants. However, it is still relatively expensive to synthesize DNA, and delivering large DNA (>50 kbp) to eukaryotic cellular compartments remains challenging. In this study, we establish a robust system for building algal mitochondrial genomes as a practical alternative to DNA synthesis. Our approach permits an inexpensive and rapid generation of mitochondrial derivatives designed for testing targeted DNA delivery. First, we cloned the mitochondrial genome of *P. tricornutum* into the eukaryotic host organism *S. cerevisiae* using two different techniques: transformation-associated recombination; and PCR-based cloning. Next, we analyzed the cloned genomes by multiplex PCR, and correct genomes were transferred to prokaryotic host *E. coli.* Then these genomes were again analyzed by multiplex PCR, followed by diagnostic digest and complete-plasmid sequencing to evaluate the fidelity of each cloning method. Finally, we assessed the burden on eukaryotic and prokaryotic hosts to propagate the cloned genomes. We conclude that our system can reliably generate variants for genome-level engineering of algal mitochondria.

**HIGHLIGHTS:**

- TAR-cloned mitochondrial genome of *P. tricornutum* in yeast
- Developed PCR-based cloning method to create designer algal mitochondrial genomes
- Stably propagated algal mitochondrial genomes in *S. cerevisiae* and *E. coli* hosts

## 1. Introduction

Pressing challenges in agriculture, medicine, and energy can be addressed through the use of designer organisms with engineered traits. The promise of synthetic biology lies in the ability to build, deliver, and install synthetic genomes and biosynthetic pathways in specialized hosts. *Phaeodactylum tricornutum* is a model diatom algal species that is an attractive candidate for synthetic biology applications [1–6]. For example, *P. tricornutum* is a popular candidate for biofuel production due to its natural propensity for lipid storage [7]. Due to the industrial and academic interest in this algal species, its genomes (nuclear, mitochondrial, and plastid) were sequenced [8–10]. The availability of genome sequences allowed for the development of genetic tools and DNA delivery methods such as biolistic-mediated transformation [11,12], electroporation [13–15] and bacterial conjugation [16,17]. Additional tools for *P. tricornutum* include a method for cloning whole chromosomes in yeast and *Escherichia coli* [18], characterized centromeres for maintaining episomal DNA [19], and genome-editing technologies [6,20–22]. We now have a powerful arsenal of tools for engineering algal nuclear genomes; however, tools for engineering and delivering organelle genomes are still lacking.

There are several advantages of engineering organelle genomes and installing synthetic DNA in these compartments rather than the nucleus. First, the polycistronic gene organization, lack of transgene silencing, and reduced positional effects in organellar DNA simplify genome engineering relative to the nucleus. Second, organelles allow for the compartmentalization of biosynthetic pathways, which isolates toxic intermediates, increases flux, and minimizes competing reactions from the rest of the cell [23]. To exploit these benefits, scientists have begun cloning whole organelle genomes from various organisms including human, mouse, maize, rice, and some algae [24–29]. Yet, a rapid method for whole organelle genome engineering for eukaryotic algae has not been established.

The first hurdle to overcome is the need for a fast and inexpensive method for synthesizing, assembling, and storing synthetic organelle genomes. While processes for synthesizing and replacing chromosomes have been established for some prokaryotes [30–34] and eukaryotes [35], genome-scale engineering and delivery is still very challenging [36]. The large, repetitive elements and AT-rich sequences of organelle genomes complicate the synthesis and cloning processes. Spurious expression of cloned genomes can be toxic to host organisms including *Saccharomyces cerevisiae* and *E. coli* [37]. Importantly, the targeted delivery of whole organelle genomes to the appropriate cellular compartment in eukaryotic algal cells is still not possible.

Here, we report a rapid protocol for cloning the *P. tricornutum* mitochondrial genome and demonstrate its maintenance in eukaryotic and prokaryotic host strains as the first step of a platform for robust, genome-scale organelle engineering. We used transformation-associated recombination (TAR) cloning to capture the wild-type mitochondrial genome of *P. tricornutum* and developed a PCR-based approach to clone modified versions of the genome. We then demonstrated the maintenance of wild type and modified mitochondrial genomes in *S. cerevisiae* and *E. coli*.

## 2. Material and Methods

### 2.1 Strains and Growth Conditions

*Phaeodactylum tricornutum* (Culture Collection of Algae and Protozoa CCAP 1055/1) was grown in synthetic seawater (liquid L1 medium) without silica at 18°C under cool white fluorescent lights (75 μE m^−2^ s^−1^) and a photoperiod of 16 h light : 8 h dark. Liquid L1 media was made as previously described by [6]. *Saccharomyces cerevisiae* VL6–48 (ATCC MYA-3666: MATα his3-Δ200 trp1-Δ1 ura3–52 lys2 ade2–1 met14 cir^0^) was grown at 30°C in rich yeast medium (2 × YPDA: 20 g yeast extract, 40 g peptone, 40 g glucose, and 200 mg adenine hemisulfate dihydrate), or synthetic complete medium lacking histidine, supplemented with adenine sulfate (60 mg L^−1^) (Teknova, Inc., Cat #: C7112). Solid yeast media contained 2% agar. After spheroplast transformation, all complete minimal media used contained 1 M sorbitol [38]. *Escherichia coli* (Epi300, Lucigen, Cat #: EC300110) was grown at 37°C in Luria Broth (LB) supplemented with chloramphenicol (15 μg mL^−1^).

### 2.2 DNA Preparation

#### 2.2.1 Isolation of *P. tricornutum* DNA in agarose plugs

For TAR-cloning, genomic DNA was isolated from *P. tricornutum* in agarose plugs. *P. tricornutum* (1.0 × 10^7^ cells mL^−1^) was plated and grown on 1%-agarose L1 plates for four days. Next, 1 mL of 1 M sorbitol was used to scrape algal cells from each plate. Next, the combined cell suspension was centrifuged at 2000 rcf for 2 min in 15 mL tube, followed by 3000 rcf for 1 min. Then, the supernatant was removed, and the cell pellet was resuspended in 2 mL of SPEM solution (1 M sorbitol, 10 mM EDTA pH 7.5, Na_2_HPO_4_ · 7H_2_O (2.08 g L^−1^), NaH_2_PO_4_ · 1H_2_O (0.32 g L^−1^)) and incubated for 1 minute at 37°C. The resuspended cells were then incubated for 5 min at 50°C and then mixed with an equal volume of 2.2% low-melting-point agarose in TAE buffer (40 mM Tris, 20 mM acetic acid, and 1 mM EDTA), which was also kept at 50°C. Aliquots of 100 μL were transferred into plug molds (BioRad, Cat #:170-3713) and allowed to solidify for 10 min at 4°C. To digest algal cell walls, the plugs were removed from the molds into 50-mL conical tubes containing 5 mL of protoplasting solution ((8.5 mL of SPEM solution, 500 μL zymolyase-20 T solution (50 mg mL^−1^), 500 μL lysozyme (25 mg mL^−1^), 500 μL hemicellulase (25 mg mL^−1^), 50 μL β-mercaptoethanol (14.3 M)) and incubated for 30 mins at 37°C. Next, the protoplasting solution was removed and the plugs were incubated with 5 mL of Proteinase K solution (100 mM EDTA (pH 8.0), 0.2% sodium deoxycholate, 1% sodium lauryl sarcosine, and 1 mg mL^−1^ Proteinase K) for 24 h at 50°C. Then, the plugs were washed four times as follows: twice with 25 mL of wash buffer (20 mM Tris and 50 mM EDTA, pH 8.0) for 2 h each at room temperature (RT), once with 25 mL of wash buffer containing 1 mM PMSF for 2 h at RT, and a final wash with 25 mL of wash buffer for 2 h.

For restriction digestion, the plugs were placed in 1.5-mL microcentrifuge tubes and washed additionally with 1 mL of 0.1X wash buffer for 1 h at RT, followed by a 5-h wash in 1X restriction buffer at RT. Finally, each plug was incubated with 50 units mL^−1^ of restriction enzyme in 1 mL of restriction buffer for 4 h at 37°C. Following the digest, the plugs were washed for 1 h in 1 mL of TE buffer (pH 8). Next, the TE buffer was removed and a fresh 100 μL of TE buffer (pH 8) was added, and plugs were melted for 10 min at 65°C. The solution was then equilibrated to 42°C for 10 min before adding 2 μL of β-agarase (NEB). Finally, the solution was incubated at 42°C for 1 h to allow the agarose to be digest. On average, each plug yielded a DNA concentration of 300 ng μL^−1^.

#### 2.2.2 DNA isolation by modified alkaline lysis

DNA from *E. coli*, yeast, and algae was isolated as described by [16]. Before isolating plasmid DNA from *E. coli* for diagnostic restriction digests or sequencing, cells were induced with arabinose. For this plasmid induction, *E. coli* overnight cultures were diluted 1:50 into 50 mL of LB media supplemented with chloramphenicol (15 μg mL−1) and arabinose (100 μg mL^−^1) and grown for 8 h at 37°C.

### 2.3 DNA preparation for PCR cloning

PCR amplification of mitochondrial fragments was performed using *P. tricornutum* genomic DNA (first iteration) or mitochondrial DNA isolated from *E. coli* as described in *Methods 2.2.2* (second iteration) as template DNA. The mitochondrial genome was amplified as eight overlapping fragments (primers: BK 141-144F/R, 145F/250R, 247F/146R, 147F/R, and 148F/140R, listed in Supplementary Table 1), as well as two additional fragments (primers: BK 88F/245R and 251F/88R, listed in Supplementary Table 1) to amplify the pAGE3.0 plasmid [17].

Each fragment was individually amplified in a 25-μL PCR reaction using 1 μL of PrimeSTAR GXL polymerase (Takara Bio Inc., #R050A), 1 μL of template DNA (10-100 ng μL^−1^ genomic DNA or plasmid DNA isolated from *E. coli*), and the respective forward and reverse primers each at a final concentration of 0.2 μM. The thermocycler was programmed as follows: five cycles of 98°C for 10 seconds, 50°C for 15 seconds, and 68°C for 480 seconds, followed by 25 cycles of 98°C for 10 seconds, 55°C for 15 seconds, and 68°C for 480 seconds, and one cycle of 68°C for 600 seconds, finishing with an infinite hold at 12°C. PCR product amplification was confirmed by performing agarose gel electrophoresis with 2 μL of PCR product on a 1.4% agarose (w/v) gel.

To eliminate plasmid template DNA, PCR products were treated with 10 units (0.5 μL) of DpnI restriction endonuclease (New England Biolabs Ltd., #R0176), incubated at 37°C for 30 minutes, and deactivated for 20 minutes at 80°C. Fragments were then purified using the EZ-10 Spin Column PCR Products Purification Kit (BioBasic Inc., #BS363) and were combined into a single 1.5-mL microcentrifuge tube to equimolar concentrations (approximately 200 ng of each fragment) and a total volume of ~30 μL.

### 2.4 DNA fragment preparation for transformation-associated recombination (TAR) cloning

*P. tricornutum* genomic DNA was digested with PvuI restriction enzyme, as described in section 2.2.1. The pAGE3.0 plasmid was amplified as two fragments (primers: BK 88F/93R & 92F/88R, listed in Supplementary Table 1). Each fragment was individually amplified in a 20 μL PCR reaction using 0.8 μL PrimeSTAR GXL polymerase (Takara Bio Inc., #R050A), 0.8 μL of template DNA (10 ng μL^−1^ of plasmid template DNA isolated from *E. coli*), and the respective forward and reverse primers at a final concentration of 0.2 μM. The thermocycler conditions used were as follows: 30 cycles of 98°C for 10 seconds, 62°C for 15 seconds, and 68°C for 150 seconds, one cycle of 68°C for 600 seconds, and an infinite hold at 12°C. PCR product amplification was confirmed by performing agarose gel electrophoresis with 1 μL of PCR product on a 1.4% agarose (w/v) gel.

To eliminate the template plasmid DNA from the PCR products, DpnI treatment, and purification was performed as described above. Then combined into a single 1.5-mL microcentrifuge tube the fragments were added in the following proportions: 10 μL linearize genome (~300 ng μL^−1^), and 7 μL for each plasmid backbone fragment (~300-500 ng μL^−1^) prior to yeast transformation.

### 2.5 Yeast Spheroplast Transformation Protocol

Yeast spheroplasts were prepared as described by [38], with the exception that mixtures of DNA fragments replaced the bacterial culture. After yeast transformation, 300 and 700 μL of yeast cell suspension were each added to a 15-mL Falcon tube containing 8 mL of melted 2%-agar yeast media lacking histidine supplemented with adenine sulfate and sorbitol, which had been kept at 50°C in a 15 mL Falcon tube. After 4–6 gentle inversions, the mixture was poured on top of an agar plate containing 10 mL of yeast media lacking histidine supplemented with adenine sulfate and sorbitol and containing 2% agar. The plate was then incubated at 30°C for 3–5 days until colonies emerged for screening.

### 2.6 *E. coli* Transformation

TransforMax Epi300 electrocompetent *E. coli* cells (Lucigen, Cat #: EC300110) were thawed on ice for 20 mins. Then, 20-μL aliquots of *E. coli* cells were transferred to sterile 1.5-mL microcentrifuge tubes and mixed with 1 μL of total DNA extracted from a single yeast colony. A Gene Pulser Xcell Electroporation System (BioRad) was set to 25-μF capacitance, 200-Ω resistance, and 2.5-kV voltage. The mixture of DNA and *E. coli* was then transferred to an ice-cold 2-mm electrocuvette and electroporated. Immediately, 1 mL of SOC media was added to the electrocuvette, which was then incubated at 37°C for 30 mins, without shaking. Next, the mixture was transferred to a sterile 1.5-mL microcentrifuge tube and incubated for 1 h at 37°C, shaking at 225 rpm. Finally, 100 μL of transformed cells were plated on selective LB media supplemented with chloramphenicol (15 μg mL^−1^) and incubated overnight at 37°C.

### 2.7 Screening Strategy

#### 2.7.1 Screening yeast colonies

To identify positive clones generated using the PCR and TAR cloning methods, all individual yeast colonies were struck onto selective 2%-agar plates lacking histidine supplemented with adenine sulfate, and grown overnight at 30°C. Next, each streak was passed onto a second selective 2% agar plate and again grown overnight at 30°C. For TAR-cloning, approximately 10 yeast colonies were pooled (20 pools in total) by picking up a small amount of each colony and resuspending them together in 100 μL of TE buffer. For PCR-based cloning, the same protocol was followed except that cells from 20 individual colonies were used for each reaction. Subsequently, for both TAR-cloning and PCR-based cloning methods, the resuspended cells were incubated at 95°C for 15 mins. The tubes were then centrifuged for approximately 30 seconds using a mini centrifuge (Scilogex, D1008). Next, 1 μL of the supernatant was used as the DNA template for diagnostic Multiplex (MPX) PCR.

MPX primer pairs were designed to have an optimized melting temperature of 60°C using the online tool Primer3 (http://bioinfo.ut.ee/primer3-0.4.0/). The MPX PCR was performed according to the Qiagen Multiplex PCR Handbook. To test potential pPT-TAR and pPT-PCR clones, two sets of MPX PCR were run, each yielding three amplicons. The first primer set, BK901, 904, and 906 F/R, was used to generate amplicons of 265-, 334-, and 507-bp sizes, respectively. The second primer set, BK 902, 903, and 905 F/R, were used to generate 171-, 224-, and 405-bp amplicons, respectively. 2 μL of the reaction products were loaded onto a 2% agarose gel for electrophoresis and then analyzed. Next, DNA was isolated (as described in 2.2.2) from selected positive clones and transformed into *E. coli* (as described in **2.6**).

#### 2.7.2 Screening *E. coli* colonies

First, one to eight *E. coli* colonies transformed with DNA from positive yeast clones were screened using the same MPX PCR method as described above for yeast colonies.

Next, the selected positive colonies were screened by diagnostic digestion. To perform diagnostic digestion, DNA was isolated from *E. coli* as described in section *Methods 2.2.2* (plasmids were induced with arabinose), and then the concentration of total isolated DNA was obtained using the DeNovix Inc. DS-11 FX+ Spectrophotometer. DNA preparations were approximately 5000 ng μL^−1^ before digestion. Digestion reactions were generated using 5 uL of DNA, 2 μL of 3.1 NEB buffer (Cat #: B7203S), 0.2 μL of SrfI (Cat #: R0629L), 0.2 μL of SacII (Cat #: R0157L), and 12.6 μL water. Reaction mixtures were incubated at 37°C for 60 minutes and then 1 μL was loaded onto a 1% agarose gel for electrophoresis.

For further confirmation, the isolated plasmid DNA was submitted to the CCIB DNA Core at Massachusetts General Hospital for whole plasmid sequencing and reference mapping. Sequences obtained were aligned with their respective references using the algorithm built into Geneious version 2020.0, created by Biomatters. Alignment disagreements were identified as mutations, and each mutation was individually analyzed to curate a list of mutations (Supplemental Table 2).

### 2.8 Evaluation of growth phenotypes of host strains

#### 2.8.1 *E. coli* growth in liquid media

*E. coli* Epi300 strains harbouring pPT-PCR C1 and C2, pPT-TAR C1 and C2, and pAGE3.0 and pPtGE31 control plasmids (lacking mitochondrial genome), were inoculated and grown overnight in 5 mL of LB media supplemented with 15 μg mL^−1^ chloramphenicol at 37°C with shaking at 225 rpm. The saturated cultures were diluted 100-fold into 5 mL of the same media and grown for 2 h in 50-mL conical Falcon tubes under the same conditions. The cultures were placed on ice and diluted to an OD_600_ of 0.1 in LB media supplemented with 15 μg mL^−1^ chloramphenicol (uninduced) or LB media supplemented with 15 μg mL^−1^ chloramphenicol and 100 ug mL^−1^ arabinose (induced). In quadruplicate, 200 μL of each uninduced and induced culture was aliquoted into a 96-well plate. Once loaded, the 96-well plate was incubated in a 96-well plate reader, Epoch 2 (BioTek). While in the plate reader, the strains were incubated at 37°C with continuous, double-orbital shaking. OD_600_ measurements were taken every 15 mins for 5 h, for a total of 20 readings using Gen5 data analysis software version 3.08 (BioTek). This experiment was performed twice, therefore for each strain, eight measurements were obtained and averaged, and the standard deviation of the mean was calculated. The doubling time (td) of each strain was determined.

#### 2.8.2 *S. cerevisiae* growth in liquid media

*S. cerevisiae VL6-48* strains harbouring pPT-PCR C1 and C2, pPT-TAR C1 and C2, and pAGE3.0 and pPtGE31 control plasmids (lacking mitochondrial genome), were inoculated and grown overnight in 5 mL of synthetic complete yeast medium lacking histidine at 30°C with shaking at 225 rpm. The saturated cultures were diluted 100-fold into 5 mL of the same media and allowed to grow for 2 h in a 50 mL conical Falcon tube under the same conditions. The cultures were diluted to an OD_600_ of 0.1 in the same media, and 200 μL of each culture was aliquoted into a 96-well plate, in quadruplicate. Once loaded, the 96-well plate was incubated in a 96-well plate reader, Epoch 2 (BioTek). While in the plate reader, the strains were incubated at 30°C with continuous shaking. OD_600_ measurements were taken every 15 mins for 12 h for a total of 49 readings using Gen5 data analysis software version 3.08 (BioTek). For each strain, four measurements were obtained and averaged, and the standard deviation of the mean was calculated. The doubling time (Td) of each strain was determined.

#### 2.8.3 *E. coli* and *S. cerevisiae* growth on solid media

*E. coli* and *S. cerevisiae* strains harbouring pPT-PCR C1 and C2, pPT-TAR C1 and C2, and pAGE3.0 and pPtGE31 control plasmids (lacking mitochondrial genome), were inoculated and grown overnight in LB media supplemented with 15 μg mL^−1^ chloramphenicol at 37°C, and synthetic complete yeast medium lacking histidine at 30°C, respectively. The saturated cultures were diluted 100-fold in their corresponding media and grown for 2 h. The cultures were diluted to an OD_600_ of 0.1, which was used to generate a range of dilutions for *E. coli* and *S. cerevisiae*. The dilution series were plated using 5 μL aliquots onto their corresponding selection plates: 1.5% agar LB plates supplemented with 15 μg mL^−1^ chloramphenicol (uninduced) or with 15 μg mL^−1^ chloramphenicol and arabinose (induced) for *E. coli*, and 2% agar synthetic complete yeast medium plates lacking histidine for *S. cerevisiae*. Dilution plates for *E. coli* strains were grown overnight at 37°C, and dilution plates for *S. cerevisiae* strains were grown for two days at 30°C.

## 3. Results and Discussion

### 3.1. Design-Build-Test cycle to enable rapid engineering of organelle genomes

Currently, there are no established methods for replacing whole eukaryotic algal organelle genomes. To address this need and enable full use of the organelle compartment for installing synthetic genomes, we are developing a design-build-test cycle for efficient capture, manipulation, delivery, and installation of organelle genomes (Figure 1A).

**Figure 1.**
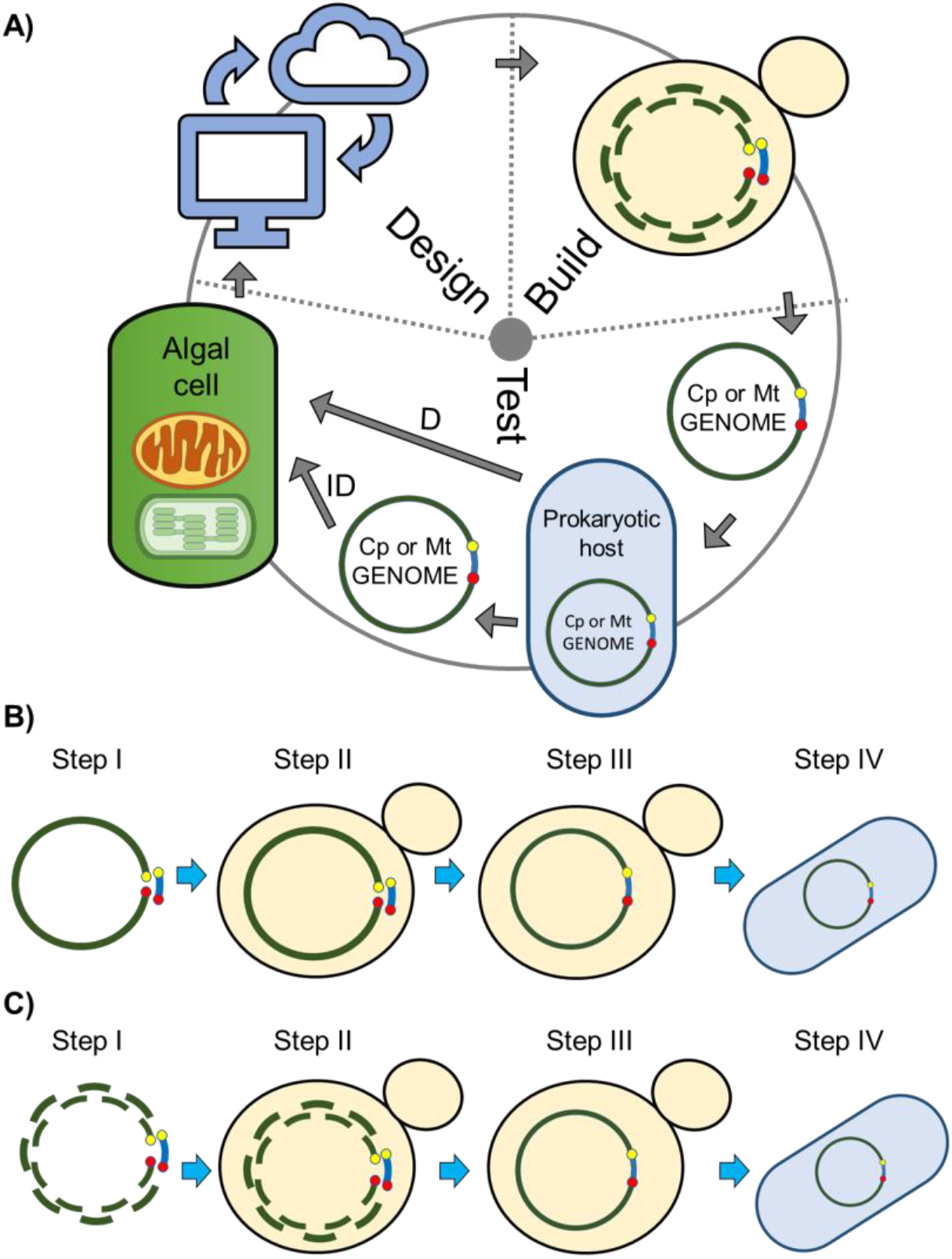
Design-Build-Test cycle for the rapid engineering of organelle genomes. **A)** Genomes will be designed based on existing knowledge and discoveries from previous cycles; in the build stage, genomes will be synthesized, then assembled and cloned in yeast; in the test stage, genomes will be isolated from yeast, moved to an intermediate prokaryotic host and delivered directly (D) (e.g. conjugation, cell fusion) or indirectly (ID) (e.g. electroporation, biolistic-mediated transformation) to the destination organelle compartment to test for viability, function, and localization. **B-C)** Schematic of the approaches used to clone the mitochondrial genomes of *P. tricornutum*. **B)** TAR cloning method; **C)** PCR-based cloning method. **Step I** differs between the PCR and TAR cloning methods where, for the PCR method, multiple overlapping fragments (green) are amplified, while for the TAR cloning method, genomic DNA is linearized at a specific location. For both methods, the plasmid backbone (blue) contains homology overlaps to the appropriate fragments or location in the genome. **Steps II-IV** are the same for both methods including DNA transformation **(Step II)** and assembly in *S. cerevisiae* via homologous recombination **(Step III)**, and transfer of clone genomes into *E. coli* **(Step IV)**.

In one iteration of the design and build stages of the cycle, we cloned a whole mitochondrial genome and a reduced version with a repetitive region removed. To capture the whole mitochondrial genome, transformation-associated recombination (TAR) cloning was used (Figure 1B). For TAR cloning, total genomic DNA was prepared in agarose plugs to obtain isolated, intact organelle genomes. Next, the mitochondrial DNA was linearized by a restriction enzyme that recognizes a single cut site in the targeted organelle genome. If there are no unique restriction enzyme cut-sites available, it is possible to select a restriction enzyme that cuts in multiple locations and perform a partial digest to obtain a proportion of genomes with a single cut at the desired location. If the cut-site resides within an essential gene, a recoded version (to remove the restriction site) of the gene can be added to the plasmid backbone. Alternatively, once the genome is cloned in yeast, the plasmid backbone can be moved to a different location by co-transformation of a plasmid containing homology hooks to the new location and another fragment that will delete the plasmid in the original location and restore the interrupted gene. Finally, a CRISPR/Cas9 system can be devised to produce appropriate cut-sites at the desired location [39,40].

The linearized algal mitochondria genome was captured by transforming it into *S. cerevisiae* along with a PCR-amplified plasmid backbone containing homology on each end to the organellar regions flanking the cut-site. Although the process is more time-consuming, an entire organellar genome can be cloned. In PCR-based cloning (Figure 1C), the mitochondrial genome was cloned indirectly by amplifying fragments from total genomic DNA with homologous DNA overhangs, followed by transformation into *S. cerevisiae* with the plasmid backbone. This process has been shown to allow for fast assembly of large plasmids [41] but it also has some drawbacks including the risk of introducing unwanted point mutations and difficulty amplifying larger repetitive elements, which could limit the practicality and versatility of this approach. In both methods, elements required for replication in yeast (e.g. *CEN6-ARSH4-HIS3*) and *E. coli* (e.g. the origin of replication, and selectable marker), must be provided as one or multiple plasmid backbone fragments containing appropriate homologous overhangs.

### 3.2. Cloning of the *P. tricornutum* mitochondrial genome

Using the PCR-based approach, we cloned a reduced version of the *P. tricornutum* mitochondrial genome (Figure 2A). The wild-type genome was PCR-amplified in eight overlapping fragments that excluded a 35 kbp region of direct repeats [9]. In place of this repeat region, two fragments with genetic elements required for replication and selection in *S. cerevisiae*, *E. coli*, and algae (nuclear localization) were amplified from the multi-host shuttle plasmid pAGE3.0 [17]. In total, 10 DNA fragments were amplified (Figure 2B) and assembled following transformation and homologous recombination in *S. cerevisiae* spheroplasts, yielding 5023 transformed yeast colonies (Table 1). After moving the assembled plasmids to *E. coli*, two clones, pPT-PCR C1 and C2, that were validated by multiplex (MPX) PCR and restriction digests (Figure 2C-D) were sequenced and analyzed for mutations (Section 3.3).

**Figure 2.**
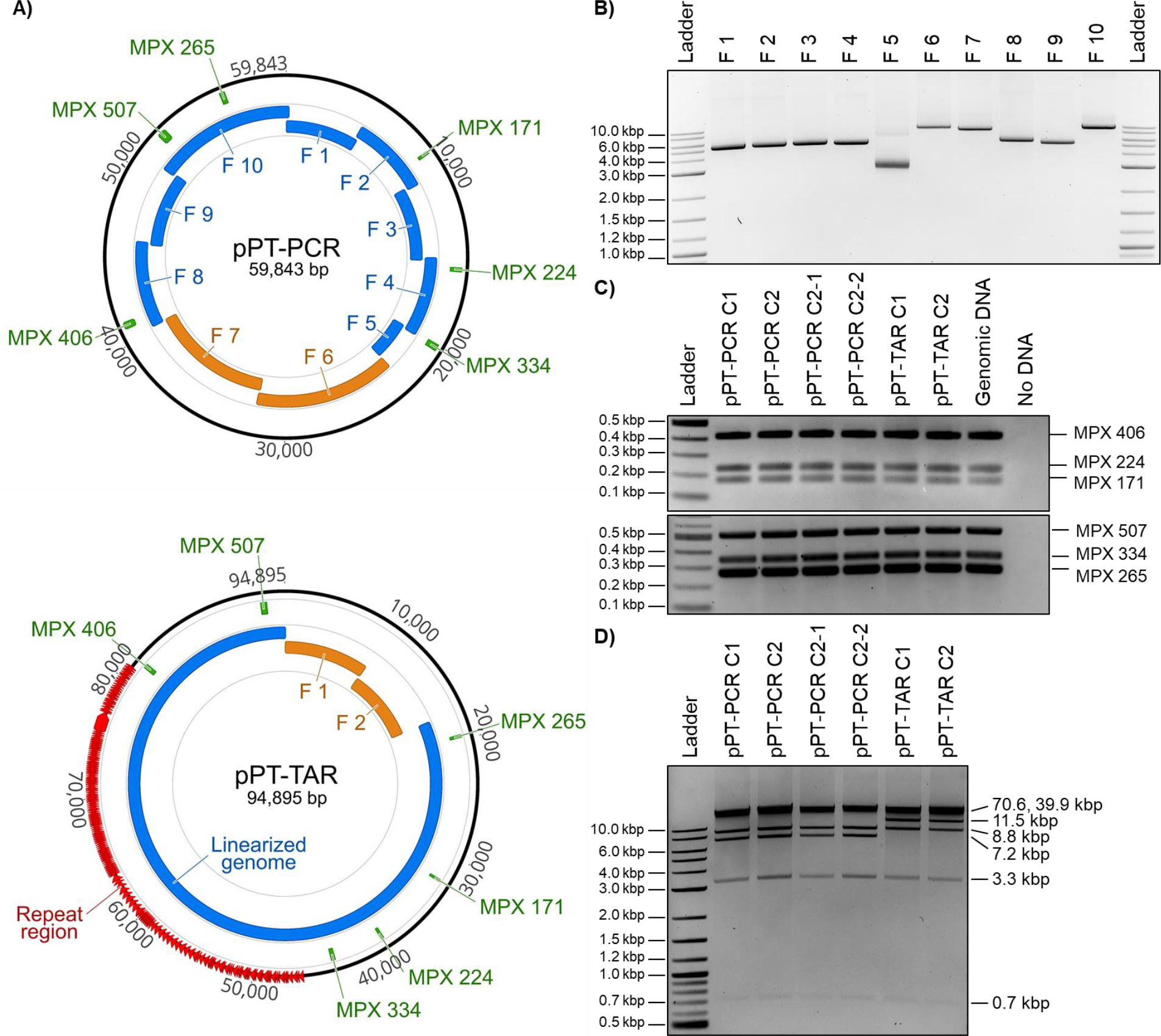
Design, amplification, and analyses of cloned *P. tricornutum* mitochondrial genomes. **A)** Plasmid maps of *P. tricornutum*’s mitochondrial genomes cloned using PCR (top) or TAR (bottom) methods. For the PCR-based cloning method, the relative sizes and positions of the eight mitochondrial fragments (blue), and the two plasmid backbone fragments (orange) are shown. For the TAR-based cloning method, the relative sizes and positions of the linearized genome (blue) and the two plasmid backbone fragments (orange) are shown. In addition, the six MPX PCR amplicons used for diagnostic screening are indicated (green). These images were generated using Geneious version 2020.0, created by Biomatters. **B)** Agarose gel electrophoresis of the 10 PCR amplified fragments of the *P. tricornutum* mitochondrial genome for PCR-based cloning including the pAGE3.0 backbone (Fragments 6 and 7). The resulting amplicon sizes for Fragments 1 to 10 are 5217, 5190, 5197, 5175, 2765, 9124, 8571, 5605, 5299, and 9264 bp, respectively. **C)** MPX PCR screen of six cloned algal mitochondrial genomes isolated from *E. coli*. The expected size for each MPX amplicon is indicated by its name (in bp). **D)** Diagnostic restriction digest of six cloned algal mitochondrial genomes using SrfI and SacII. For the pPT-PCR clones, the expected band sizes are 39925, 8758, 7177, 3276, and 707 bp; and for the pPT-TAR clones, the expected band sizes are 70647, 11506, 8758, 3276, and 707 bp. Notes: 1) We did not see a size difference between the 70.6-kbp and 39.9-kbp fragments, which is most likely due to the electrophoresis conditions (1% agarose gel, 100 V for 90 minutes); 2) The 707-bp band is present but very faint. For all gels, we used NEB 2-log ladder.

**Table 1.** Cloning of the *P. tricornutum* mitochondrial genome in the host organisms *S. cerevisiae* and *E. coli.* Two PCR-cloning assemblies and one TAR-cloning assembly were performed. For the *E. coli* selection media, CM indicates chloramphenicol antibiotic.

| Design<br>Assembly Type<br>DNA Source | *S. cerevisiae* |  |  | *E. coli* |  |  |  |
| --- | --- | --- | --- | --- | --- | --- | --- |
|  | Selection<br>Media | Colony<br>Count | MPX PCR<br>Screen<br>Positive/Total | Selection<br>Media | Selected Yeast<br>Colony: *E. coli*<br>Colony Count | MPX PCR<br>Screen<br>Positive/Total | Final Genomes<br>Selected For<br>Analysis |
| Reduced Genome<br>PCR - 10 Fragments<br>Genomic DNA | -Histidine | 5023 | 20/20 | CM | Yeast colony C1:30<br>Yeast colony C2: 18 | C1=8/8<br>C2=8/8 | pPT-PCR C1<br>pPT-PCR C2 |
| Reduced Genome<br>PCR - 10 Fragments<br>Clone pPT-PCR C2 | -Histidine | 4880 | 20/20 | CM | N/D | C2-1=5/5<br>C2-2=5/5 | pPT-PCR C2-1<br>pPT-PCR C2-2 |
| Full Genome<br>TAR - 3 Fragments<br>Digested Genomic DNA | -Histidine | 604 | 2/204 | CM | Yeast colony C1: 119<br>Yeast colonyC2: 46 | C1=1/1<br>C2=2/2 | pPT-TAR C1<br>pPT-TAR C2 |

Importantly, we wanted to assess the fidelity of the PCR-based cloning method by comparing cloned sequences to their parental reference. However, the amplified fragments initially used in this iteration were derived from a heterogenic population of mitochondrial genomes, which may differ from the published sequence. Therefore, to directly determine the mutation rate of the cloning method, the mitochondrial genome was re-amplified using DNA isolated from an isogenic culture of *E. coli* carrying pPT-PCR C2. The 10 fragments were assembled using the same method, with similar results as the first assembly (Table 1, Section 3.3). Two additional clones, pPT-PCR C2-1 and C2-2, were selected for further analysis.

To clone the complete, repeat-containing genome, we used the TAR-based approach (Figure 2A). Total *P. tricornutum* genomic DNA was digested with the restriction enzyme PvuI, which cuts the mitochondrial genome at a single site. We amplified the plasmid backbone from pAGE3.0 as two fragments with end homology to each other and the genomic sequences flanking the PvuI cut site. Transforming both PCR-amplified plasmid backbone fragments and the linearized genome into *S. cerevisiae* spheroplasts yielded 204 colonies (Table 1) from which two clones, pPT-TAR C1 and C2, were identified as positive for the presence of the mitochondrial genome using MPX PCR. We then transferred these genomes to *E. coli* and analyzed again with MPX PCR and performed a diagnostic restriction digest (Figure 2C-D). Although the build stage using the TAR-cloning approach was successful, it was time-consuming and labor-intensive due to the requirement of obtaining high-quality DNA and screening a larger number of colonies. Nonetheless, this iteration can act as a good template for future design-build cycles, particularly for cloning large organellar genomes that are difficult to PCR-amplify.

### 3.3. Sequence analysis of cloned *P. tricornutum* mitochondrial genomes

The six selected *P. tricornutum* mitochondrial clones were sequenced and analyzed for mutations. Sequences obtained for the original PCR-cloned (pPT-PCR C1 and pPT-PCR C2) and TAR-cloned (pPT-TAR C1 and pPT-TAR C2) plasmids were aligned to reference sequences based on the published sequence from Oudot-Le Secq and Green (2011) and the plasmid from Brumwell et al. (2019); and the reassembled PCR-cloned plasmids (pPT-PCR C2-1 and pPT-PCR C2-2) were aligned to the sequence of their parent clone, pPT-PCR C2. Upon analyzing mutations in pPT-PCR C1 and C2, an average of 7 mutations per 60 kbp was found (Table 2), which corresponds to 1 mutation per 8.6 kbp. The two reassembled clones, pPT-PCR C2-1 and C2-2, acquired an average of 3 mutations per 60 kbp (Table 2), corresponding to 1 mutation per 20 kbp. See Supplementary Table 2 for additional information about specific mutations. pPT-TAR C1 had only a single synonymous substitution (Table 2). However, the other clone generated using this method, pPT-TAR C2, contained a series of deletions in the repetitive region of the *P. tricornutum* mitochondrial genome (Table 2). This clone also carried seven base-pair substitutions and an insertion of 43 bps, albeit all in noncoding regions of the plasmid. The difference in mutation rate between these two plasmids could be the result of recombination of the highly repetitive region of the mitochondrial genome during TAR cloning, or an artifact of the sequencing method used as resolving repetitive regions may require the use of long-read sequencing technologies.

**Table 2.** Summary of mutations identified in the cloned *P. tricornutum* mitochondrial plasmids. Identified mutations are categorized as point mutations (synonymous, missense, nonsense, and those found in non-coding regions) or gap mutations (insertions and deletions, either noncoding or resulting in frameshifts).

| Clone | Point Mutations |  |  |  | Gap Mutations |  |  |  | Total |
| --- | --- | --- | --- | --- | --- | --- | --- | --- | --- |
|  | Synon-<br>ymous | Missense | Nonsense | Non-<br>coding | Noncoding |  | Deletions |  |  |
|  |  |  |  |  | Insertions | Deletions | Insertions | Deletions |  |
| pPT-PCR C1 | 0 | 4 | 0 | 2 | 1 | 1 | 1 | 0 | 9 |
| pPT-PCR C2 | 0 | 3 | 1 | 0 | 0 | 1 | 0 | 0 | 5 |
| pPT-PCR C2-1 | 0 | 3 | 0 | 2 | 0 | 0 | 0 | 0 | 5 |
| pPT-PCR C2-2 | 0 | 1 | 0 | 0 | 0 | 0 | 0 | 0 | 1 |
| pPT-TAR C1 | 1 | 0 | 0 | 0 | 0 | 0 | 0 | 0 | 1 |
| pPT-TAR C2 | 0 | 0 | 0 | 7 | 1 | 8 | 0 | 0 | 19 |

Genetic changes found using both cloning methods could have occurred during the cloning process of each mitochondrial plasmid or during propagation in the host organisms harboring the plasmids. It is also plausible that some of these variants could naturally exist in the heterogeneous population of *P. tricornutum* mitochondrial genomes. If desired, identified mutations could be fixed through additional iterations of the design-build-test cycle.

### 3.4. Maintenance of *P. tricornutum*’s mitochondrial plasmids in host organisms

*S. cerevisiae* and *E. coli* were used as host organisms to capture and store *P. tricornutum*’s mitochondrial genome. *S. cerevisiae* was chosen because it is currently the best organism for assembling large DNA molecules due to its highly efficient homologous recombination machinery and demonstrated ability to maintain a wide range of genomes without adverse effects [42]. However, when using standard protocols for isolating plasmid DNA, the yields from *S. cerevisiae* tend to be low. To overcome this problem, after assembly in *S. cerevisiae*, the cloned mitochondrial plasmids were transformed into *E. coli*. In *E. coli*, the mitochondrial plasmids, which contain an arabinose-inducible origin of replication, can be induced to a higher copy number to generate increased DNA yields. Importantly, propagating genomes in *E. coli* allows for the development of direct transfer methods, such as conjugation, to deliver the plasmids to the desired destination organism. At the same time, because mitochondrial genomes are of prokaryotic origin, there is a chance that a prokaryotic host would express functional proteins that might have adverse or toxic effects.

We sought to examine the burden of propagating the cloned mitochondrial genomes in eukaryotic and prokaryotic host strains. Colony formation on selective plates and growth rates in liquid media for both *S. cerevisiae* and *E. coli* strains carrying these plasmids were evaluated. If a plasmid causes cellular stress, it can decrease the growth rate and result in the formation of smaller or fewer colonies on the plate. In *S. cerevisiae*, there was no observable decrease in colony size when strains harbouring cloned genomes were spot plated, as compared with control plasmids that have the same backbone, pAGE3.0 and pPtGE31^6^ (Figure 3A). Next, we compared growth in liquid media by using a 96-well plate reader. Interestingly, the strains carrying the mitochondrial genomes grew faster in liquid as compared to the pAGE3.0 control, but at a similar rate when compared to the pPtGE31 control (Figure 3C). Overall, this indicates that the maintenance of *P. tricornutum*’s mitochondrial genomes plasmids does not have an adverse effect on yeast growth. In *E. coli*, we did not observe any substantial negative effects on growth rates of cells harbouring *P. tricorunutm’s* mitochondrial genomes, as compared to the control plasmids (Figure 3B and D). In addition, there was no substantial effect on growth rates when the cells were induced to have high copy number of mitochondrial genomes.

**Figure 3.**
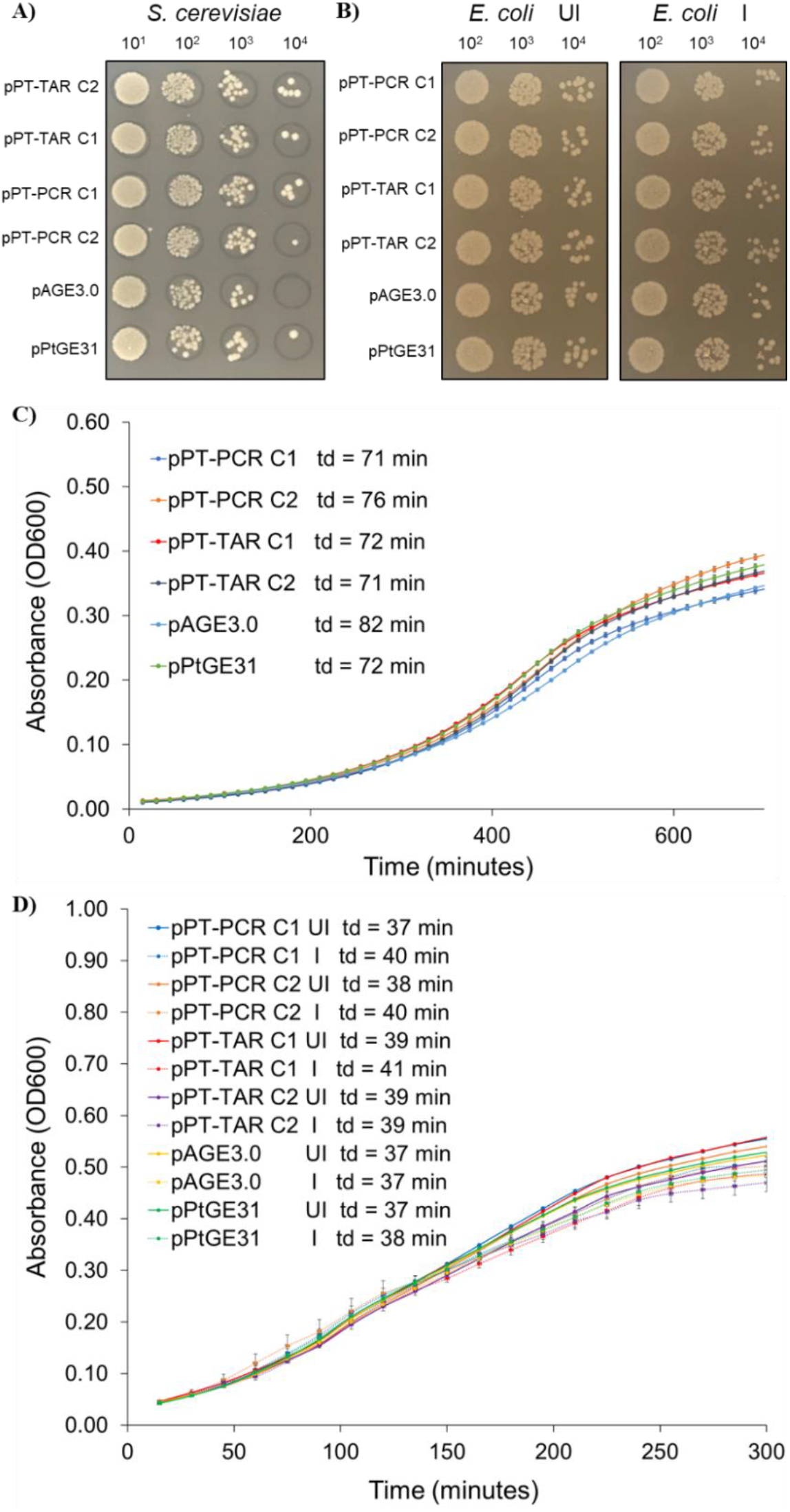
Analysis of growth of *S. cerevisiae* and *E. coli* strains harbouring cloned *P. tricornutum* mitochondrial genomes on solid media and in liquid media. **A)** Dilutions (10^1^-10^4^) of *S. cerevisiae* strains plated on solid synthetic complete media lacking histidine. **B)** Dilutions (10^2^-10^4^) of *E. coli* strains plated on solid LB media supplemented with chloramphenicol only (UI - uninduced) or with chloramphenicol and arabinose (I - induced) to increase plasmid copy number. **C)** Growth curves of *S. cerevisiae* strains grown in liquid synthetic complete media lacking histidine. **D**) Growth curves of *E. coli* grown in liquid LB media supplemented with chloramphenicol only (UI) or with chloramphenicol and arabinose (I). Td: doubling time.

## 4. Conclusions

The biotechnological potential of organelle engineering is held back by the lack of reliable methods to replace organellar genomes. Towards the goal of establishing a design-build-test cycle for genome-scale organelle engineering, we have developed two adaptable methods for cloning and manipulating eukaryotic mitochondrial genomes. With a PCR-based approach, we cloned a variant of the *P. tricornutum* mitochondrial genome lacking its 35-kbp repeat region and with a TAR-based approach, we captured the complete genome. The former had a mutation rate ranging from 1 mutation per 8.6-20 kbp, while the latter allowed us to identify one clone with only a single mutation. The cloned genomes imposed no substantial growth burden on *S. cerevisiae* and *E. coli* when these host organisms were used to propagate the plasmids. In this study, we completed the first step in developing a reproducible set of methods for cloning, manipulating, and installing synthetic organelle genomes.

## Acknowledgements

This work was supported by Natural Sciences and Engineering Research Council of Canada (RGPIN-2018-06172) awarded to B.J.K. Salary for P.J. was supported by Designer Microbes Inc. (DMI). In addition, research in D.R.E., is also supported by Natural Sciences and Engineering Research Council of Canada (NSERC), RGPIN-2015-04800.

## Authorship Contributions

**Ryan R. Cochrane:** Formal analysis, Investigation, Methodology, Validation, Visualization, Roles/Writing - original draft; Writing - review & editing.; **Stephanie L. Brumwell** : Formal analysis, Investigation, Validation, Visualization, Roles/Writing - original draft, Writing - review & editing.; **Maximillian P.M. Soltysiak** : Investigation, Validation, Writing - review & editing.; **Samir Hamadache** : Formal analysis, Investigation, Validation, Visualization, Writing - review & editing.; **Jennifer G. Davis** : Investigation.; **Jiayi Wang** : Investigation.; **Samuel Q. Tholl** : Investigation.; **Preetam Janakirama** : Investigation.; **David R. Edgell** : Resources, Supervision, Writing - review & editing.; **Bogumil J. Karas:** Conceptualization, Funding acquisition, Investigation, Methodology, Project administration, Resources, Supervision, Validation, Visualization, Roles/Writing - original draft, Writing - review & editing.

## Appendix A. Supplementary Data

Supplementary data to this article can be found online

