## Supplemental Information for "Rapid method for generating designer algal mitochondrial genomes"

**Supplementary Table 1.** List of primers used in the amplification and screening of *Phaeodactylum tricornutum*'s mitochondrial genome.

| **Name** | **Primers** | **Length (bp)** |
|---|---|---|
| | **Amplification of *P. tricornutum*'s Mitochondrial Genome: pPT-PCR** | |
| Fragment 1 | **BK141F –** ctgttacacgttggactgggacaaaatggtttattgaagg<br>**BK141R –** ttggaataaaagggttcgaacctttgaatgatcgtaccaa | **5217** |
| Fragment 2 | **BK142F –** tttatttaacaaaatcgcctcttggggttcttatgcatca<br>**BK142R –** actagtcgttaaatggttagttagtttgtttttaataagc | **5190** |
| Fragment 3 | **BK143F –** tggtcgggtaaaagacctacctggagtaaaatatcatcta<br>**BK143R –** gttttcccaaagatggggggcgatgttttttccatttta | **5197** |
| Fragment 4 | **BK144F –** gtgtaaatctatgaaaaagttattgaaatcaaaattacga<br>**BK144R –** tccatgttttattaaaaatatttgaatgtttcagtatttt | **5175** |
| Fragment 5 | **BK145F –** cccagtttcatttttgggctccaatctaaatgtcgttaaa<br>**BK250R –** cgacaagcacgagcgagatatcccaatcaagctagtatcgatcgcgcgcgggcgaagcttccctaaaaagatgacaaacc | **2815** |
| Fragment 6 | **BK251F –** tttctgtaaatcggtttgtcatctttttagggaagcttcgcccgcgcgcgatcgatactagcttgattgggatatctcgc<br>**BK88R –** aagcttgaccgagagcaatcccgcagtcttcagtggtgtgatggtcgtctatgtgtaagtcaccaatgcactcaacgatt | **9124** |
| Fragment 7 | **BK88F –** aatcgttgagtgcattggtgacttacacatagacgaccatcacaccactgaagactgcgggattgctctcggtcaagctt<br>**BK245R –** tcccccccgcctaaggaaacttggtcgtaaaagcgtgaaaagcgatcgcgatcgtcttgccttgctcgtcggtgatgta | **8571** |
| Fragment 8 | **BK247F –** tcgagctgtaagtacatcaccgacgagcaaggcaagacgatcgcgatcgcttttcacgcttttacgaccaagtttcctta<br>**BK146R –** cttcacggaaaattcaatttcgctgagtttgtgctggaga | **5655** |
| Fragment 9 | **BK147F –** gctgacgcctgcccagtgctgcaagattaaaggaaagggt<br>**BK147R –** agtaagcacggcgaaaaaaaaggtagaactggtaggagat | **5299** |
| Fragment 10 | **BK148F –** tagcttttcgctccgaaacccaagatgtttttttttcatc<br>**BK140R –** cccaacactattaaattcttcaactttgtttacaggattt | **9264** |
| | **TAR Cloning Capture Vector: pPT-TAR** | |
| Fragment 1 | **BK92R –** ttgcatttttttgggcgttctttcattctagtatagcacctgcccatacgatggcgcgccatcgatactagcttgattgg<br>**BK88R –** aagcttgaccgagagcaatcccgcagtcttcagtggtgtgatggtcgtctatgtgtaagtcaccaatgcactcaacgatt | **9142** |
| Fragment 2 | **BK88F –** aatcgttgagtgcattggtgacttacacatagacgaccatcacaccactgaagactgcgggattgctctcggtcaagctt<br>**BK93F –** gacagtgaaggtctttataaggcttattgctttaggcggccaaactcctccgcgtcgacggatcgtcttgccttgctcgt | **8581** |
| | **Diagnostic Multiplex PCR Primers** | |
| Amplicon 1 | **BK901F –** tattgcatcgaggcacagag<br>**BK901R –** gcccaaagcataggtgtcat | **265** |
| Amplicon 2 | **BK902F –** aaagctgcaaggcagttgat<br>**BK902R –** aggccaaaaaggtttcgatt | **171** |
| Amplicon 3 | **BK903F –** ggcagaaaagctgagcctaa<br>**BK903R –** cctatggttgcaaggcattt | **224** |
| Amplicon 4 | **BK904F –** ttcatttttgggctccaatc<br>**BK904R –** cagtttcaggttcgggatgt | **334** |
| Amplicon 5 | **BK905F –** gttcttgtttcgccggatta<br>**BK905R –** aacacagaccgacaccttcc | **405** |
| Amplicon 6 | **BK906F –** acgttttgcagtacccttgg<br>**BK906R –** accataagtgcacgggaatc | **507** |

**Supplementary Table 2.** List of mutations identified in cloned *P. tricornutum* mitochondrial genomes. Plasmids containing the cloned genomes were completely sequenced using an Illumina® MiSeq™ and mapped to their respective reference sequences by the CCIB DNA Core Facility at Massachusetts General Hospital (Cambridge, MA). Mutations were identified using Geneious version 2020.0 created by Biomatters. Positions are based on residue numbering beginning with the first base in Fragment 1 of the result sequence and counting towards Fragment 2 (see Figure 2A). Published Sequence: expected sequence as designed by combining published sequence for the *P. tricornutum* mitochondrial genome [1] and based on plasmid pAGE3.0 [2] according to the respective cloning strategy.

| Position | Mutation | Location/Gene | Effect |
|---|---|---|---|
| **pPT-PCR C1**<br>Reference: Published Sequence | | | |
| 8,002 | G > T | *mt-Rps2* | Arg114 > Ile |
| 15,327 | C > A | *mt-Rpl2* | Gln35 > Lys |
| 16,259 | 1-bp insertion | *mt-Rps19* | Frameshift |
| 18,073 | C > A | *mt-atp9* | Leu42 > Met |
| 22,687 | G > A | *mt-Cob* terminator | Unknown |
| 22,714 – 22,871 | 158-bp insertion | *mt-Cob* terminator | Unknown |
| 34,783 | 129-bp deletion | Vector backbone (intergenic) | No effect |
| 57,408 | G > T | Intergenic | No effect |
| 59,856 | C > T | *mt-Cox1* | Thr245 > Ile |
| **pPT-PCR C2**<br>Reference: Published Sequence | | | |
| 159 | 1-bp deletion | *mt-Cox1* intron | Unknown |
| 17,214 | G > T | *mt-Rps3* | Met310 > Ile |
| 27,716 | G > T | Vector backbone (*S. meliloti repA2*) | Ala135 > Glu |
| 42,554 | G > T | *mt-Nad5* | Glu595 > STOP |
| 56,559 | T > C | *mt-Nad7* | Ile191 > Thr |
| **pPT-PCR C2-1**<br>Reference: pPT-PCR C2 | | | |
| 1,197 | C > T | *mt-Cox1* intron | Unknown |
| 1,473 | C > T | *mt-Cox1* intron | Unknown |
| 12,811 | G > A | *mt-Nad1* | Asp295 > Asn |
| 27,716 | T > G | Vector backbone (*S. meliloti repA2*) | Glu135 > Ala |
| 55,673 | G > A | *mt-Cox2* | Val195 > Ile |
| **pPT-PCR C2-2**<br>Reference: pPT-PCR C2 | | | |
| 27,716 | T > G | Vector backbone (*S. meliloti repA2*) | Glu135 > Ala |
| **pPT-TAR C1**<br>Reference: Published Sequence | | | |

| | | | |
|---|---|---|---|
| 17,871 | C > A | *mt-Cox3* | Gly215 > Gly |
| **pPT-TAR C2**<br>Reference: Published Sequence | | | |
| 8,415 | C > A | Vector backbone<br>(*S. cerevisiae ARSH4*) | No effect |
| 12,257 – 12,299 | 43-bp insertion | Vector backbone<br>(intergenic) | Unknown |
| 17,906 | C > A | *mt-Cox3* | Gly215 > Gly |
| 46,055 | 3,847-bp deletion | Repeat region | Effect unknown.<br><br>Note: Mutations detected in repetitive region are likely due to sequencing errors. |
| 46,182 | 8,004-bp deletion | Repeat region | |
| 46,235 | T > C | Repeat region | |
| 46,246 | 403-bp deletion | Repeat region | |
| 46,300 | A > G | Repeat region | |
| 46,304 | C > T | Repeat region | |
| 46,442 | G > A | Repeat region | |
| 46,536 | G > A | Repeat region | |
| 46,851 | 8,165-bp deletion | Repeat region | |
| 46,927 | 7,906-bp deletion | Repeat region | |
| 47,185 | 497-bp deletion | Repeat region | |
| 47,757 | 3,601-bp deletion | Repeat region | |
| 47,891 | C > T | Repeat region | |
| 47,913 | 413-bp deletion | Repeat region | |
